# Widespread introgression of the *Mus musculus musculus* Y chromosome in Central Europe

**DOI:** 10.1101/2019.12.23.887471

**Authors:** Miloš Macholán, Stuart J. E. Baird, Alena Fornůsková, Iva Martincová, Pavel Rubík, Ľudovít Ďureje, Emanuel Heitlinger, Jaroslav Piálek

## Abstract

According to Haldane’s rule, sex chromosomes should harbour more incompatibilities than autosomes. As a consequence, transmission of sex-linked genes across a genetic barrier is expected to be hampered. A remarkable example of a contradiction of this assumption was reported from the hybrid zone between two house mouse subspecies in western Czechia and south-eastern Germany where unidirectional east→west Y chromosome introgression was observed. Since the phenomenon was coupled with differences in sex ratio, this was hypothesised to be caused by a genetic conflict between sex-specific genes on sex chromosomes or elsewhere in the genome. Here we capitalise on a large material consisting of almost 7500 mice collected across a vast area from the Baltic Sea to the Alps embracing a ~900 km long portion of the zone with the aim to (i) detect its exact course and (ii) reveal the extent and pattern of the Y chromosome introgression in Central Europe. We show that the path of the zone is quite tortuous even at the global scale and the introgression is rather a rule than an exception. We also show that although sex ratio perturbations described in our previous study appear also in other introgression areas, they may not be ubiquitous. Finally, we reveal that although not all Y chromosome types are associated with the introgression, it is not restricted to a single ‘winning’ haplotype.

## 1 INTRODUCTION

More than three decades ago Barton and Hewitt, in their seminal review of hybrid zones (Barton & Hewitt, 1985), concluded that the majority of hybrid zones are likely to be tension zones. According to theory, tension zones are maintained by a balance between dispersal and intrinsic selection against hybrids (Key, 1968). This has two consequences. First, differential population pressure on either side of the zone minimises its length and second, since selection against hybrids is independent of any particular habitat, the tension zone is free to move until being trapped by a geographical barrier or in an area of low population density (a ‘density trough’; Barton, 1979; Hewitt, 1975, 1989). This opens, on the other hand, room for geographic features to affect the zone course on a local scale. Hybrid zones can be viewed as semi-permeable or, better, selectively permeable, boundaries between differentiated genomes across which alleles thriving on hybrid background move quickly whereas introgression of mutations reducing hybrid fitness or engendering conspecific mating preferences is strongly counterselected. A third group of variants is equally fit in hybrids and parental populations. These alleles are expected to diffuse stochastically across the zone in both directions (Barton, 1979; Barton & Bengtsson, 1986; Barton & Gale, 1993; Barton & Hewitt, 1985; Bazykin, 1969; Harrison, 1986; Piálek & Barton, 1997). Restricted introgression is expected especially in sex-linked loci, in agreement with the ‘two rules of speciation’ (Coyne & Orr, 1989). The first is Haldane’s rule stating that when hybrids of one sex suffer from impaired viability or fertility, it is usually the heterogametic sex (Haldane, 1922), and the second one is the Large X-effect pointing to the fact that hybrid incompatibilities often map to the X chromosome (Charlesworth, Coyne, & Barton, 1987; Coyne, 1992; Coyne & Orr, 1989).

One of the best studied hybrid zones is the zone of secondary contact between two house mouse subspecies in Europe, eastern *Mus musculus musculus* (*Mmm*) and western *M. m. domesticus* (*Mmd*). This zone is more than 2500 km long and runs from Scandinavia through Central Europe to the Black Sea coast (Figure 1a). Following the pioneering works of Degerbøl (1949) and Ursin (1952) from Jutland and Zimmermann (1949) from Central Europe, the European house mouse hybrid zone (HMHZ) has been studied in a number of areas (for a recent review see Baird & Macholán, 2012). The HMHZ is a mixture of late-generation hybrids and backcrosses with F1 hybrids being either missing or extremely rare. For most markers its structure is unimodal, with intermediate genotypes and lowest fitness values in the centre (Albrechtová et al., 2012; Baird et al., 2012; Macholán et al., 2007; Raufaste et al., 2005). The zone width varies among markers but in general it is lower for X chromosome than for autosomal markers (Dod et al. 1993; Macholán et al. 2007; Tucker, Sage, Warner, Wilson, & Eicher, 1992), with most incompatibilities localised in the central X (Dufková, Macholán, & Piálek, 2011; Janoušek et al., 2012; Macholán et al., 2011; Payseur, Krenz, & Nachman, 2004; Teeter et al., 2010).

**Figure 1.**
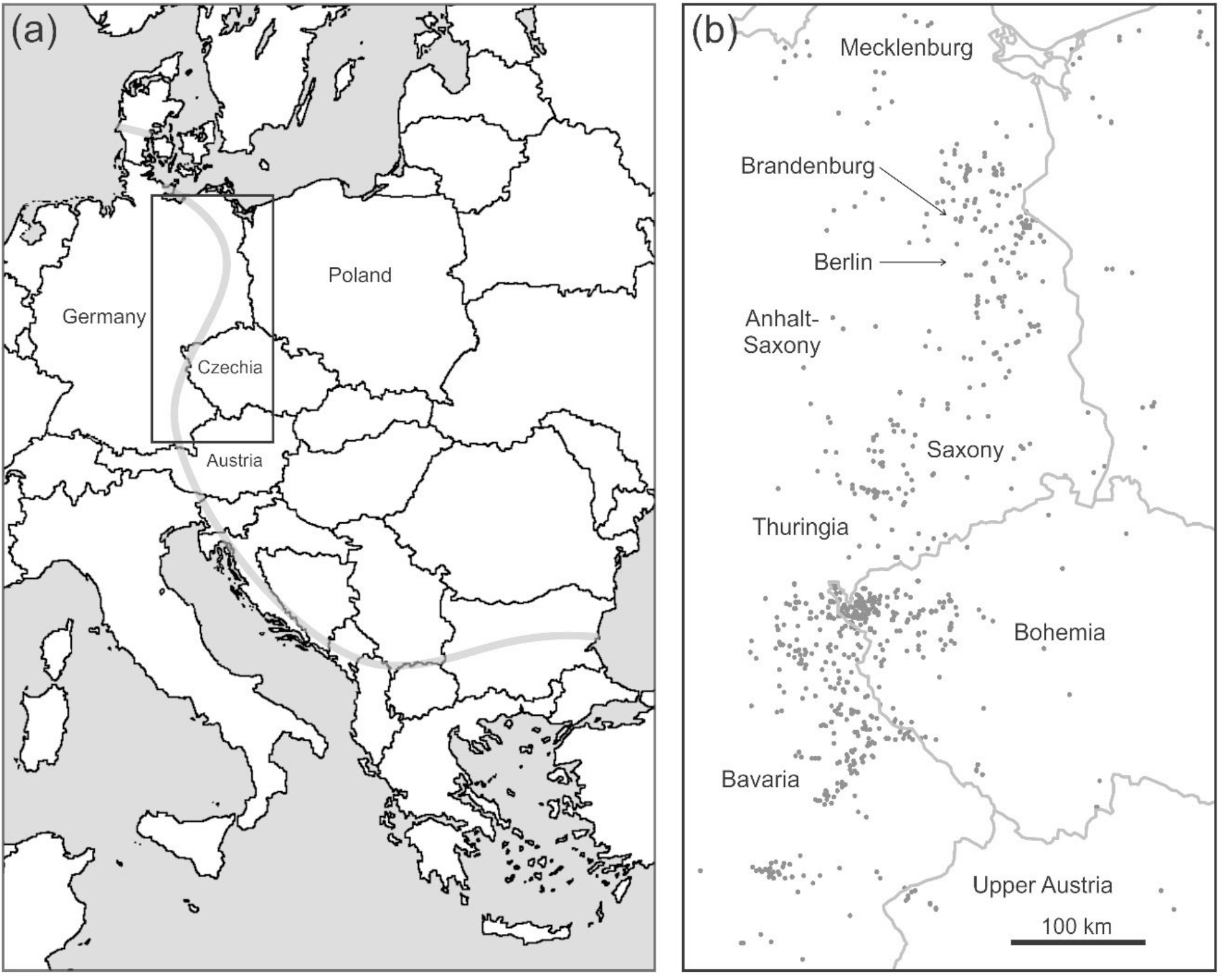
(a) A sketch of the HMHZ course in Europe (grey curve); the study area is depicted with black rectangle; (b) detail of the area with sampling localities.

Based on Haldane’s rule, Y chromosomes are expected to be strongly counterselected on alien genetic background. Reports of the absence of Y introgression across the HMHZ from Denmark (Dod, Smadja, Karn, & Boursot, 2005; Vanlerberghe, Dod, Boursot, Bellis, & Bonhomme, 1986), Bulgaria (Vanlerberghe et al., 1986), and southern Bavaria (Tucker et al., 1992) seem to be consistent with this expectation. However, Munclinger, Božíková, Šugerková, Piálek, and Macholán (2002) found *Mmm* Y chromosomes deep in the *Mmd* range in western Bohemia (Czech Republic) and north-eastern Bavaria (Germany) and this finding was later confirmed on a much larger data set (Macholán et al., 2008). These authors also revealed this phenomenon to be accompanied by differences in the census sex ratio (SR): this was significantly female-biased in the *Mmm* territory and in *Mmd* localities without the introgressed *Mmm* Y, whereas in the *Mmd* area with the introgressed *Mmm* Y it was not significantly different from parity so that there was a clear and significant difference between ‘introgressed’ *vs*. ‘non-introgressed’ regions. This led Macholán et al. (2008) to hypothesise that introgression of *Mmm* Y chromosomes is a consequence of a genetic conflict between the X and Y (with some autosomal elements likely also involved). According to this hypothesis, the *Mmm* Y is thriving on the naïve *Mmd* background (or, if the Y introgression had been preceded by another, enabling, factor, the *Mmm* Y thriving on the altered *Mmd* background) and hence spreads at the expense of the *Mmd* Y. Though often viewed rather as an exception than a rule (Jones & Searle, 2015), a recent study of Ďureje, Macholán, Baird, and Piálek (2012) has suggested that the Y introgression may not be limited to the Czech-Bavarian portion of the HMHZ. (Presence of the *Mmm* Y in western Norway far behind the zone in *Mmd* territory, reported by Jones, Van Der Kooij, Solheim, & Searle [2010], does not seem to be consistent with movement of the chromosome across the zone.)

The fact that the HMHZ course is far from being straight across Europe (Figure 1a) calls for explanation. However, for such a task we first need to detect the exact course of the HMHZ which is, strikingly, largely unknown despite a half a century of research. This is the first aim of this study. The second goal is to detect the Y chromosome introgression pattern across a vast area embracing a substantial portion of the HMHZ from the Baltic Sea to the border between southern Bavaria and North Tyrol. We show that (i) the HMHZ across the study area is rather complicated even at the global scale; (ii) the unidirectional introgression of the *Mmm* Y chromosome into *Mmd* territory is widespread in Central Europe; (iii) sex ratio perturbations previously found to be coupled with the Y introgression in the Czech-Bavarian portion of the HMHZ may not be universal; and (iv) although not all Y chromosome types are associated with the introgression, it is not restricted to a single ‘winning’ haplotype.

## 2 MATERIAL AND METHODS

### 2.1 Sampling

This study covers a 266,539 km^2^ area (377 km × 707 km) embracing an ~900 km long portion of the HMHZ running from the Baltic Sea coast through former East Germany and western Bohemia (Czech Republic) to the northern slopes of the Alps in southern Bavaria and Lower Austria (Figure 1b). The total sample consisted of 7440 mice (3522 males and 3918 females). Genetic analyses were carried out in 7270 mice (3431 males and 3839 females) captured at 804 localities. As a Y chromosome marker, we used an 18 bp deletion in the *Zfy2* gene located in the non-recombining region of the chromosome (*N* = 3286) which is fixed in *Mmm* and absent in *Mmd* (Nagamine et al., 1992; Orth et al., 1996). For comparison and to precisely detect the HMHZ course across the study area, we scored up to 44 autosomal and X-linked markers (*N* = 7270). Since these diagnostic loci are usually used for calculating some sort of hybrid index (HI) we refer to them as ‘HI markers’.

To describe and depict the Y chromosome introgression pattern we employed the spatially explicit Bayesian model-based clustering software Geneland v. 4.0.6 (Guillot, Mortier, & Estoup, 2005). We simplified the analysis by limiting the number of clusters to *K* = 2, i.e. the number of subspecies present in the study area. At least three independent MCMC runs with 5,000,000 iterations were carried out for individual genotypes. In all cases, the first 25% iterations were discarded as burn-in. The spatial model included no uncertainty ascribed to geographic coordinates and no null alleles were allowed. Both correlated and uncorrelated allele frequency models were applied; however, the results were invariant to this aspect of the model.

A previous study of the Czech-Bavarian portion of the HMHZ indicated introgression of the *Mmm* Y is accompanied by differences in the census sex ratio (Macholán et al., 2008). In the current study we define an ‘introgressed population’ in two ways. First, we follow a rather extreme position considering as introgressed each *Mmd* population sample (i.e. of HI less than 0.5) harbouring at least one *Mmm* Y chromosome. The second approach is based on combination of the curves of the Geneland-estimated centre based on the HI markers and that of the Y chromosome (i.e. areas assigned to *Mmd* with the HI markers and to *Mmm* based on the Y marker). In this way we delimit three regions: (i) the ‘Dy_D_ region’, defined either as all areas with prevalence of *Mmd* alleles and no *Mmm* Y present or as all areas assigned with Geneland to *Mmd* based on both the HI and Y markers; (ii) the ‘M region’, i.e. all areas with prevalence of *Mmm* alleles (HI > 0.5) or those assigned with Geneland to *Mmm* based on both types of markers; and (iii) the ‘DY_M_ region’ defined as described above. Within these regions we calculated sex ratios. The total sample used for this task comprised 7396 individuals (3497 males, 3899 females). The exact binomial test was used to detect deviations of census sex ratios from parity. The tests were performed in R statistical environment (R Core Team, 2018). Variances of sampling sizes within regions and hybrid zone partitions were tested with Kruskal-Wallis test using Statistica (TIBCO Software Inc., 2018).

### 2.2 *Sry* sequencing

In total, we sequenced 113 males. We amplified a 1,277-bp fragment upstream of the HMG box of the *Sry* gene (Gubbay et al., 1990) including a part of the inverted repeated flanking region (Gubbay et al., 1992), using primers F6685 and R7920 published in Nachman and Aquadro (1994). DNA was amplified in 35 cycles following a 15-min pre-heating stage at 95 °C. Each cycle consisted of 30 seconds at 94 °C, 1.5 min at 57 °C, and 2 min at 72 °C. The total volume per sample contained 5 μl of 2x Qiagen master mix, 0.3 μl of each primer (10 mM), 2.4 μl of H_2_O, and 2 μl of DNA.

We removed the (CA)_*n*_ microsatellite from the data since ascertaining accurate sequence of this highly variable repeat appeared unreliable. Hence, after including the outgroup sequences the final aligned sequences were 1,055 bp long. Phylogenetic trees were inferred using MEGA 7.0.26 (Kumar, Stecher, & Tamura, 2016). For the maximum likelihood analysis, we employed the HKY+Γ model (Hasegawa, Kishino, & Yano, 1985) selected with jModelTest2 v2.1.6 (Darriba, Taboada, Doallo, & Posada, 2012; Guindon & Gascuel, 2003) under AICc (Hurvich & Tsai, 1989; Sugiura, 1978), BIC (Schwarz, 1978), and decision theory selection (Minin, Abdo, Joyce, & Sullivan, 2003). The CIPRES Science Gateway (Miller, Pfeiffer, & Schwartz, 2010) was employed for running jModelTest program. The continuous gamma distribution was discretised to six and 10 categories for the whole data set and distinct haplotypes, respectively. Bootstrap support (Felsenstein, 1985) was based on 1000 pseudoreplicates. To improve searching for the best tree, an extensive subtree pruning and regrafting procedure (Felsenstein, 2004) was applied. The weakest stringency of optimisation with respect to branch lengths and improvements in log likelihood values (branch swap filter) was used to maximise likelihood search.

### 2.3 Microsatellites

1017 males were scored for 12 microsatellite loci based on method of Hardouin et al. (2010) and P. Rubík (pers. comm.). However, locus Y23 (Hardouin et al. 2010) was found to be located on Chromosome 1 according to the NCBI Primer-BLAST and hence it was excluded from the analysis, reducing the number of loci to 11. Forward primers were labelled with fluorescent labels on 5’ ends (primer sequences and type of labelling available in Supplementary Table S1). DNA amplification was performed using the Qiagen Multiplex PCR kit. All reactions were carried out in 5 μl final volumes using 10 ng of DNA template. Microsatellites were grouped in two multiplexes. The first multiplex contained loci Y21, Y22 and Y24 and was run under the following PCR conditions: 95 °C for 15 minutes, followed by 30 cycles of 95 °C for 30 s, 60 °C for 90 s, and 72 °C for 60s, with the final extension at 72°C for 10 min. The second multiplex consisted of loci 5035, 5045, 7245, 6132, 7322, 7419, Y6 and Y12. The PCR conditions were: 95 °C for 15 min followed by 30 cycles of 95 °C for 30 s, 51 °C for 90 s, 72 °C for 30 s and the final extension at 60°C for 30 min. Fragment sizes were genotyped using 3130x/Genetic Analyzer (Applied Biosystems) and manually scored using GeneMapper v. 3.7 (Applied Biosystems).

Since all the scored loci are located in the nonrecombining region of the Y chromosome and hence they are not independent, common approaches such as ordination analysis (principal component or coordinate analysis, multidimensional scaling etc.) or clustering using STRUCTURE or similar programs are inappropriate for our data. Another problem with these methods is the absence of any model of allele relatedness. For example, alleles 100 and 102 are likely to be more related than alleles 100 and 120 yet all are treated as completely independent by both PCA-like and STRUCTURE-like programs.

Therefore, we adopted an alternative approach to processing the microsatellite data in which we treat each Y chromosome as a single haplotype: hence, for example, the {80,96,82,100,120,108} haplotype is considered one step different from the {80,96,82,100,120,110} haplotype. For all 1017 Y chromosomes we found 353 unique encodings (haplotypes). The commonest Y haplotype was shared by 55 males. In the following step we computed all pairwise Euclidean distances between the haplotypes scaled relative to the variance of each microsatellite and taking into account structure. The distances not exceeding a threshold value, *dY*, then represent edges joining sets of vertices (haplotypes). For *dY* = 0 each unique Y haplotype forms its own cluster. As *dY* increases, a haplotype is allowed to cluster with ‘similar’ haplotypes (e.g., ones that differ by one repeat at one locus). As *dY* further increases, more and more distant haplotypes are allowed to cluster together and thus the number of distinct clusters drops, eventually leading to over-merging. The microsatellite data were processed using Mathematica v. 11.3 (Wolfram Research Inc., 2018).

## 3 RESULTS

### 3.1 The course of the HMHZ

Although the global pattern of the HMHZ course across the area under study seems to roughly coincide with the pattern inferred in previous studies (Figure 1a), zooming in points to substantial differences. The first deviation appears south of the 52^nd^ degree of latitude (north of the 5700th *y* coordinate in Figure 1) where the zone curves westwards to bend sharply southwards around the 51^st^ parallel (south of 5700) reaching the slopes of the Ore Mountains (Krušné hory in Czech, Erzgebirge in German) on the border between north-western Bohemia and southern Saxony (Figures 1b and 2a). The second departure from the global pattern appears on the border between Bohemia and north-eastern Bavaria where the zone first follows the Upper Palatine Forest (Český les/Oberpfälzer Wald) ridge up to its south-eastern margin and then makes an almost right-angle turn to the south-west (Figure 2a). As a result, instead of a smooth, more-or-less linear course between an area east of Berlin in the north-east and Munich in the south-west, the zone makes two elbow-like deflections, one westward and one eastward.

**Figure 2.**
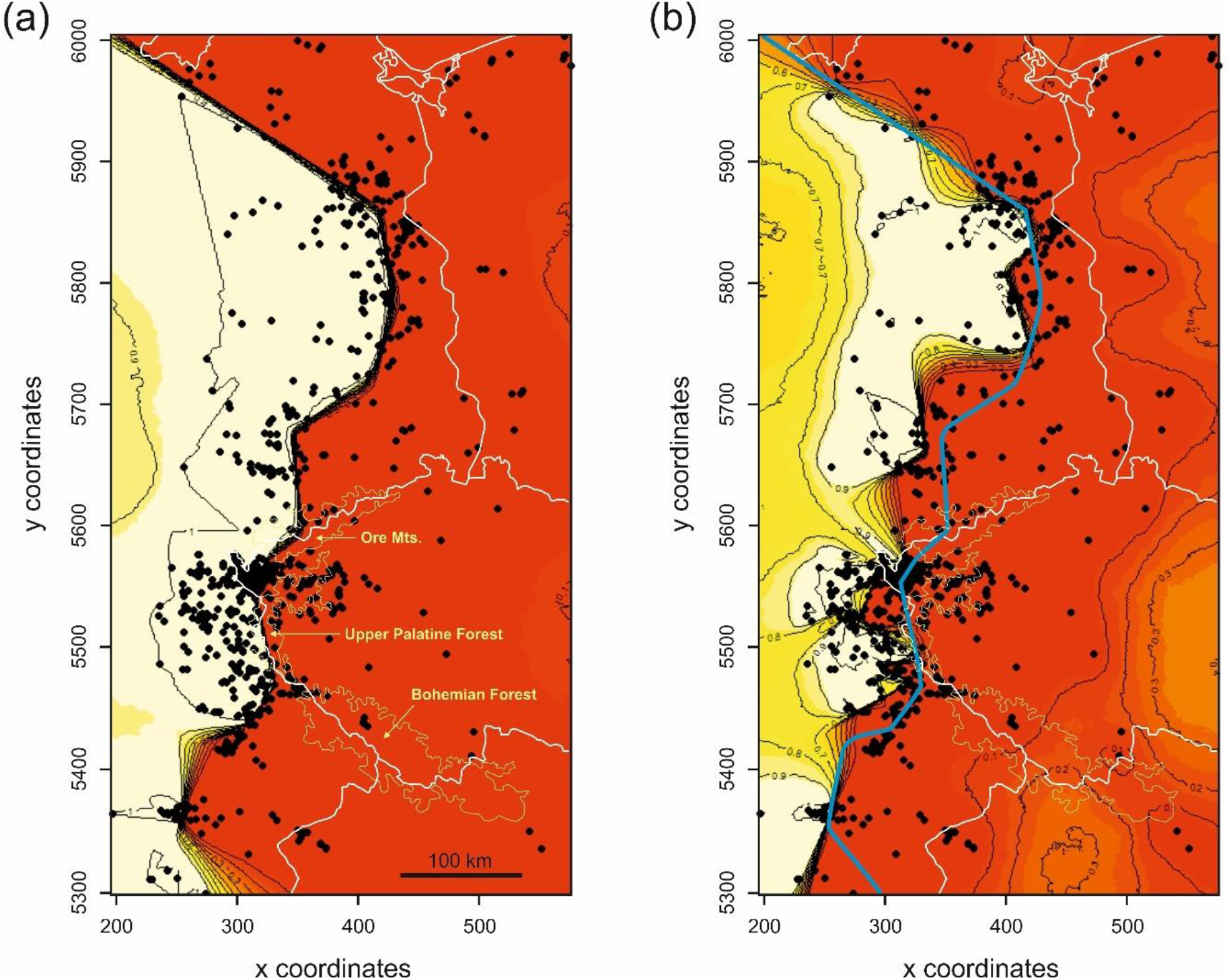
Map of the probability of each individual’s membership either to *M. m. musculus* (red colour) or to *M. m. domesticus* (light yellow) based on genotypes at the HI markers (a) and Y chromosomes (b). The level contours show the spatial changes in assignment values. For comparison, the course of the zone inferred from the HI markers is shown as bold blue line.

### 3.2 Y chromosome introgression

As shown in Figure 2b, introgression of *Mmm* Y chromosomes into *Mmd* territory is overwhelming. Comparison with the HMHZ course inferred with the HI markers reveals vast *Mmd* areas with introgressed *Mmm* Ys, the largest being in Saxony, Thuringia, and Saxony-Anhalt (the central part of Figure 2b; cf. also Figure 1b) and west of the Czech-Bavarian border, respectively. In the former area, for example, as many as 26 *Mmm* Y chromosomes, contrasting with only a single *Mmd* Y, were found in Gniebitz near Trossin (Nordsachser District), about 36 km west of the zone centre, and one *Mmm* Y appeared even 39 km west of the centre (Starkenberg, Altenburger Land Distr.). A similar degree of introgression was revealed within the Czech-Bavarian part of the study area: the most distant site with introgressed *Mmm* Y chromosomes was Plössen 18 (Bayreuth Distr.), a locality as far as 49 km west of the zone centre, where all seven genotyped males possessed the *Mmm* Ys.

There is an interesting area of ambiguity just south of the Czech-Bavarian introgreesion region (i.e., roughly south of the 5500th *y* coordinate in Figure 2b). This area is a mosaic of pure *Mmd* localities interspersed with polymorphic sites with both types of Y chromosome. This suggests a different, more diffuse and stochastic, introgression pattern.

### 3.3 Sex ratio

As expected, the more stringent definition of the DY_M_ region as a group of *Mmd* localities with at least a single introgressed *Mmm* Y made this category to embrace higher numbers of individuals than that based on the Geneland-based estimate. Nevertheless, both methods revealed similar results pointing to female-biased SR in the non-introgressed regions (DY_D_, M) contrasting with more balanced SR in the DY_M_ region (Geneland-based results not shown). While the census sex ratio was significantly female-skewed in the DY_D_ and M regions the DY_M_ region was not significantly different from parity (Table 1).

**Table 1.**
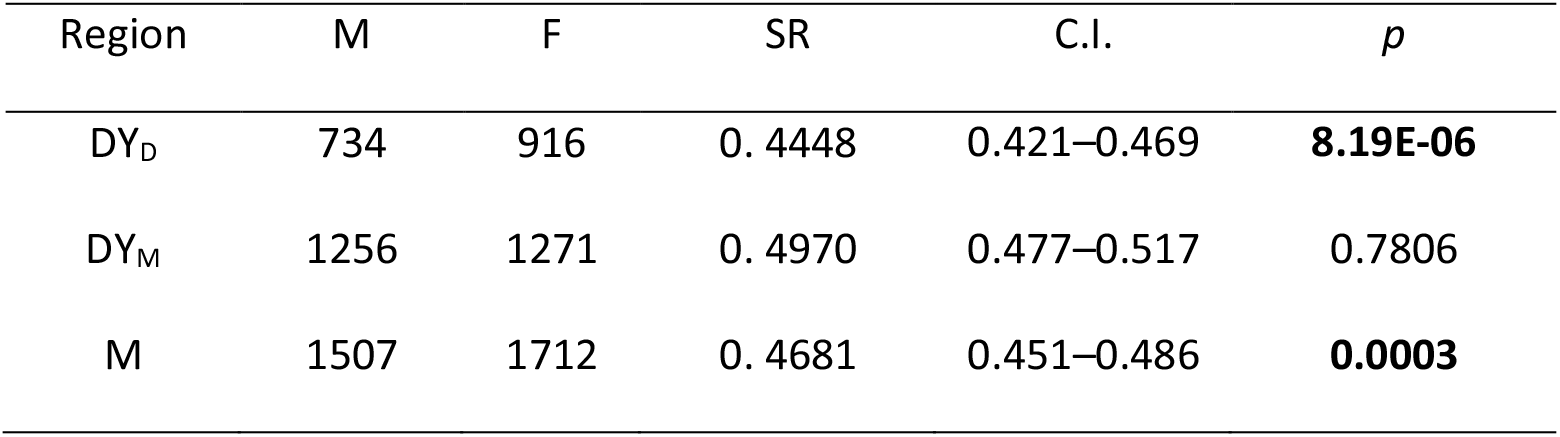
Census sex ratios (SR), expressed as proportion of males, in the whole data set. M, F = number of males and females, respectively; C.I. = 95% confidence intervals. As DY_M_ we consider any population of predominately *Mmd* background, i.e. with hybrid index HI < 0.5, harbouring at least one *Mmm* Y chromosome (see Material and Methods for details on the definition of all three regions).

| Region | M | F | SR | C.I. | *p* |
| --- | --- | --- | --- | --- | --- |
| $DY_D$ | 734 | 916 | 0.4448 | 0.421–0.469 | **8.19E-06** |
| $DY_M$ | 1256 | 1271 | 0.4970 | 0.477–0.517 | 0.7806 |
| M | 1507 | 1712 | 0.4681 | 0.451–0.486 | **0.0003** |

However, it could be argued that the data are dominated by the large sample collected from the Czech-Bavarian portion of the HMHZ which represents more than 50% of all individuals under study and so the results can be biased. Therefore, we partitioned the data to the ‘Czech’ and ‘non-Czech’ parts; the latter was then divided into northern and southern part. In this way we divided the total data set to three parts: ‘Northern’, ‘Central’, and ‘Southern’. Since the sample sizes dropped substantially due to this partitioning, especially in the ‘Northern’ and ‘Southern’ part, the deviations from parity in the DY_D_ and DY_M_, regions appeared insignificant except of that in the DY_D_ region in the ‘Southern’ partition; nevertheless, the trend remained the same as in the whole data set, the only exception being the ‘Southern’ *Mmm* region with higher number of *males* than females (Table 2).

**Table 2.** Census sex ratios (SR) in three partitions: ‘Northern’, ‘Central’, and ‘Southern’ (see Material and Methods); M and F = number of males and females, respectively; C.I. = 95% confidence intervals.

| Area | Region | M | F | SR | C.I. | *P* |
| --- | --- | --- | --- | --- | --- | --- |
| Northern | DY<sub>D</sub> | 205 | 241 | 0.4596 | 0.413–0.507 | 0.0974 |
|  | DY<sub>M</sub> | 264 | 273 | 0.4916 | 0.449–0.535 | 0.7300 |
|  | M | 390 | 443 | 0.4783 | 0.444–0.512 | 0.2177 |
| Central | DY<sub>D</sub> | 425 | 534 | 0.4432 | 0.411–0.475 | **0.0005** |
|  | DY<sub>M</sub> | 737 | 738 | 0.4997 | 0.474–0.525 | 1.0000 |
|  | M | 869 | 1059 | 0.4507 | 0.428–0.473 | **0.0000** |
| Southern | DY<sub>D</sub> | 104 | 141 | 0.4245 | 0.362–0.489 | **0.0213** |
|  | DY<sub>M</sub> | 255 | 260 | 0.4951 | 0.451–0.539 | 0.8601 |
|  | M | 248 | 210 | 0.5415 | 0.497–0.588 | 0.0837 |

Another source of bias can be sample size. We can, for example, expect males, as the more dispersing sex in house mice, to dominate smaller samples. Indeed, we found the proportion of males to be significantly negatively correlated with sample size (*R*^2^ = 0.0050; *p* = 0.0455). However, no significant differences in sample sizes were revealed among the regions within each partition (Kruskal-Wallis test: *p* > 0.05 for all three partitions). We can thus conclude that varying sample sizes are unlikely to be responsible for the differences in SR between the regions.

### 3.4 Sequencing

In general, the sequences were highly similar: in all 113 sequences there were as few as 22 distinct haplotypes (7_ *Mmm*, 12_*Mmd*, 3_*M*. *macedonicus* used as outgroup). The two subspecies were only distinguished by two diagnostic positions, all other substitutions were either singletons or rare alleles shared by two or a few sequences. As a consequence, there was no internal substructuring within the subspecies.

As shown in the maximum likelihood tree (Supplementary Figure S1), the males form two clusters, *Mmm* and *Mmd*. As expected, the *Mmm* clade also includes the predominantly *Mmd*-derived classical laboratory strains A/J, C57BL, and C3H known to bear *Mmm*-type Y chromosome and males of predominately *Mmd* background carrying introgressed *Mmm* Ys. Both clusters are highly homogeneous, revealing no internal phylogenetic structure.

### 3.5 Microsatellites

When all 353 unique haplotypes are allowed to merge into two clusters the resulting groups more or less correspond to Y chromosome types discriminated with the diagnostic *Zfy2* gene: only ~1% of haplotypes assigned to the ‘eastern’ group had *Mmd* Ys and ~2% of haplotypes assigned to the ‘western’ group had *Mmm* Y chromosomes (data not shown). With decreasing *dY* more and more clusters appear, the process starting within *Mmm* territory. Figure 3 depicts geographic distribution of 12 haplotype groups. In *Mmd*, there are two main groups, one restricted to the northern and western parts of the area under study (‘blue haplotypes’) and the other occurring in the central and southern parts (‘cyan haplotypes’). Other haplotypes are locally restricted. In *Mmm*, there are three dominant groups. While the most widespread ‘red’ and a bit less common ‘orange’ haplotypes are introgressing far to the south in *Mmd* territory, the ‘yellow’ haplotypes are, with few exceptions, restricted to the invasion salient stretching between the northerly Ore Mts. and the southerly Upper Palatine Forest (cf. Figure 2a). Again, other haplotypes are either locally restricted or do not show a clear geographic pattern (Figure 3). We can thus conclude that the *Mmm* introgression is not restricted to a single ‘winning’ haplotype.

**Figure 3.**
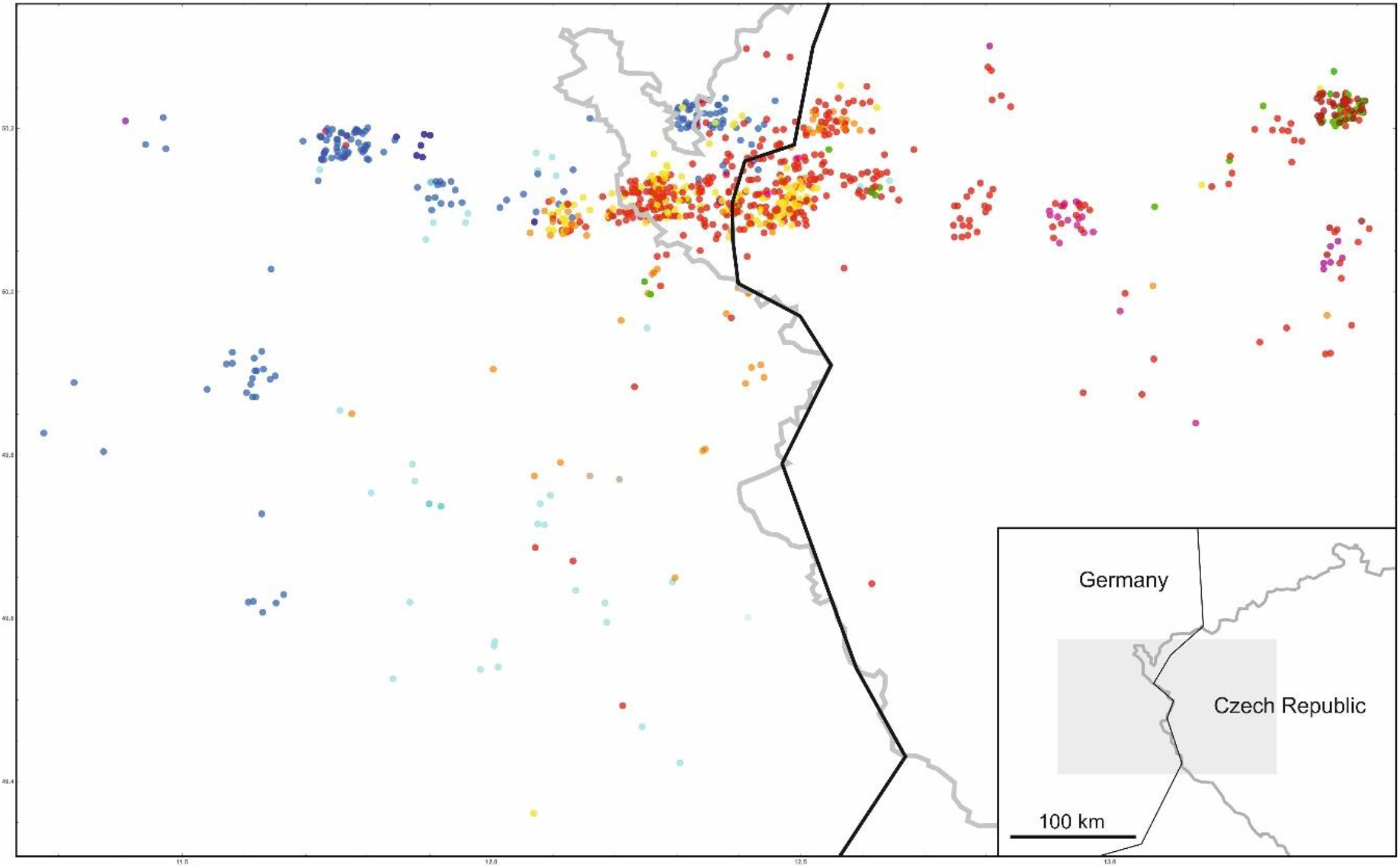
The border area between western Bohemia, Czech Republic, and north-eastern Bavaria, Germany (insert); large figure depicts position of Y chromosomes clustered into 12 haplogroups. Grey lines = state borders; black lines = hybrid zone centre.

## 4 DISCUSSION

### 4.1 HMHZ course

As introduced above, tension zones should tend to minimise their lengths (Barton 1979; Barton and Hewitt 1985; Key 1968). So why the HMHZ does not follow a straight line across Europe? One possible explanation is that the zone is affected also by extrinsic selective forces. For example, ranges of the two mouse subspecies can be influenced by climatic conditions, namely by the boundary between oceanic and continental climate zones (Boursot et al. 1984; Hunt & Selander, 1973; Klein, Tichy, & Figueroa, 1987; Serafiński, 1965; Thaler, Bonhomme, & Britton-Davidian, 1981; Zimmermann, 1949). However, the scale of the climatic transition seems to be too wide to maintain such a narrow hybrid zone (Boursot, Auffray, Britton-Davidian, & Bonhomme, 1993). In addition, the two gradients, ecological and genetical, are not coincident, as pointed out already by Kraft (1985) and further discussed by Baird and Macholán (2012). Hence although we cannot *a priori* rule out a potential influence of some climatic factors on house mouse populations, especially in northern areas, in the absence of convincing evidence we should search for other causes shaping the HMHZ course in Europe.

Another theoretical prediction is that inasmuch as tension zones are not located at a particular environmental gradient, they can move until being trapped by a geographical barrier or in an area of low population density (Barton, 1979; Hewitt, 1975, 1989). Hence the current position of the zone could reflect either presence of physical barriers, a consequence of historical contingencies or simply represents a transitional stage. All three possibilities may contribute to the zone dynamics on a global scale and even (intricately interweaved) on a local scale. In the context of the present study the most obvious physical barriers are forested mountain ridges along the border between Germany and Czech Republic. Larger forests are known to hamper house mouse dispersal (Zejda, 1975). This barrier was further strengthened by a drastic decrease of human population densities in these areas after the World War II and sealing off the frontier after 1948. In other parts of the HMHZ the factors affecting its position are less clear and will be analysed elsewhere.

### 4.2 *M. musculus musculus* Y chromosome introgression

Here we showed that, in downright contradiction to theoretical expectations, introgression of the *Mmm* Y chromosomes into the *Mmd* range is almost ubiquitous (at least) in Central Europe. Given the magnitude of this phenomenon it is interesting to look at areas with *no* introgression. One apparent example seems to be the area north of Munich (bottom left part of Figure 2b). Indeed, absence of any Y chromosome introgression in this region was reported by Tucker et al. (1992). This could be explained by the barrier effect of the Moosach River (not the ‘floodplains of the Isar River’ suggested by Sage, Whitney, and Wilson [1986]) and the vast forest area west of Freising. However, a closer look reveals a scatter of previously studied localities (Sage et al. 1986; Teeter et al. 2008; Tucker et al. 1992) with *Mmm* Y chromosomes introgressing up to about 7 km into the *Mmd* range (Ebertspoint, Geselthausen, Giggenhausen, Massenhausen, Neufahrn, Thalhausen). Moreover, at more recently sampled sites west and south of Munich the *Mmm* Y introgression was found to be even deeper. So, the only regions with no Y chromosome introgression appear in northern parts of the study area, most notably those south of Berlin (ca. 52^nd^ parallel; coordinate 5750 in Figure 2b) and north of Berlin (ca. 53^rd^ parallel; coordinate 5850 in Figure 2b).

However remarkable this may seem, the glaring differences in the degree of *Mmm* Y introgression are, in fact, not astounding. We may hypothesise that either different *Mmm* Y chromosomes encounter the same *Mmd* genetic background in different portions of the HMHZ or vice versa (we cannot rule out a third possibility of meeting both different *Mmm* Y’s and different *Mmd* backgrounds either). Yet the discordant introgression pattern may not require genetic differences between local consubspecific populations. If a genetic element (notwithstanding if selfish or unselfish) is advantageous on the alien genetic background, upon secondary contact it is predicted to quickly introgress across the hybrid zone and spread far away from it (Barton & Bengtsson, 1986; Barton & Gale, 1993; Barton & Hewitt, 1985; Piálek & Barton, 1997). However, if these loci are tightly linked to barrier genes, they may be unable to quickly recombine or segregate away from their background. Then it is just a matter of time when these ‘ticking bombs’ disengage themselves from linkage and launch their introgressive spread. And because recombination is random introgression is not expected to be homogeneous in space along the whole zone of contact. Instead, advancing alleles may form spatial irregularities like the salients we see in Figure 2b (Ibrahim, Nichols, & Hewitt, 1996; Macholán et al., 2008, 2011).

The role of geographic features as barriers to Y chromosome introgression is most conspicuous in the hilly and forested frontier area between western Bohemia and north-eastern Bavaria (Upper Palatine Forest Mts.). Due to this barrier *Mmm* Y chromosomes can only intrude into the area west of these mountains either from the south or the north or from both directions. While Figure 3 appears to reject the first possibility in favour of the second, according to the Geneland-based results (Figure 2b) *Mmm* Ys introgress from both directions. The microsatellite data point to an interesting difference in the *Mmm* Y introgression pattern in north-eastern Bavaria (Figure 3): while the transition is very steep in the northern part, between the Ore and Upper Palatine Forest Mts. (see also Macholán et al. 2008), it is rather diffuse southwards. This contrast appears to be coincident with the distribution of two widespread *Mmd* Y chromosome types (blue vs. cyan haplotypes in Figure 3).

## AUTHOR CONTRIBUTION

The study was designed by MM and JP; the data were gathered by MM, AF, IM, PR, ĽĎ, EH, and PKT, and analysed by MM, SJEB, IM, and JP. mM, SJEB, JP, and IM contributed to writing the paper.

## Notes

FUNDING INFORMATION, This work was supported by the Czech Science Foundation (grant No. 15-13265S to MM and SJEB).

