## Supplementary Material for "Widespread introgression of the *Mus musculus musculus* Y chromosome in Central Europe"

**Table S1.** The list of Y chromosome microsatellites with primer sequences and labelling dye.

| NAME | F/R | Sequence 5’ - 3’ | Label | Length | Source |
| --- | --- | --- | --- | --- | --- |
| Y6 | F  R | AACCACCACTATCTTCATTC  ACAGAGTATACGTACGTGTG | VIC | 120-124 | Hardouin et al. (2010) |
| Y12 | F  R | CCCAATCTAGGCATTTAATT  ATTCACCATTCTCCAGTGTG | 6FAM | 118-140 | Hardouin et al. (2010) |
| Y21 | F  R | TCATGGTAGACACCATGGCAAC  TCAGTTTTCTAGGTGGAGGGGTG | PET | 239-294 | Hardouin et al. (2010) |
| Y22 | F  R | ACCATCAGATGATCACCAAGTGC  TCCAGCATTCAATGGTACAGGCT | NED | 295-324 | Hardouin et al. (2010) |
| Y24 | F  R | TCTGGGGGTTTCGGGTGGAGCCT  GCATCACAGCTGAGGCTCTGTGG | 6FAM | 373-399 | Hardouin et al. (2010) |
| 5036 | F  R | TGCTGTGGAAGTGAACAGAAA  AACTAGCCAGGTCACCAGACA | 6FAM | 74-86 | Rubík (2011)^a^ |
| 5045 | F  R | TGAAATAAACCAGGGCAAAAA  ACATGATTGCTAACCCCTTCC | PET | 109-124 | Rubík (2011) |
| 6132 | F  R | AAAGAAGAGCCAGGAGTGAGC  TTACCCAGCTGTTCTTCCCTT | NED | 140-166 | Rubík (2011) |
| 7245 | F  R | GGGTCCTTAAAATTGGTTGCT  AGTAAAGGAGGCCGATCATGT | PET | 143-166 | Rubík (2011) |
| 7322 | F  R | TGACCACTCTGGCTCATCTTT  CATGAGCTAATTTGCCTCTGC | NED | 154-184 | Rubík (2011) |
| 7419 | F  R | CGTTCCAATATCAACCCCTTT  CTGTTGCAAAAGCAAAAGACA | NED | 188-227 | Rubík (2011) |

^a^ Unpublished diploma thesis: Rubík, S. (2011). Y chromosome in the house mouse hybrid zone. Charles University in Prague (in Czech with English abstract).

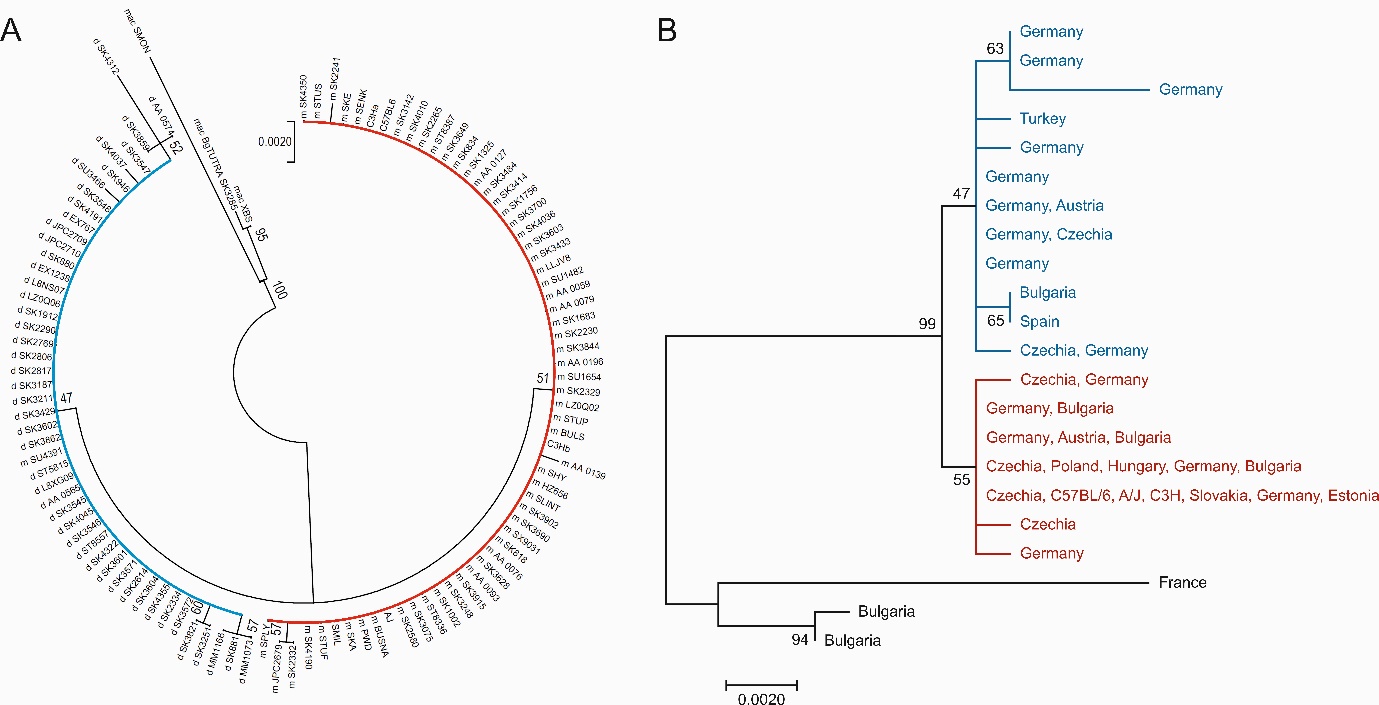

**Figure S1.** Maximum-likelihood tree inferred with *Sry* sequences; (A) full data set; (B) distinct haplotypes only. Three *M. macedonicus* sequences were used as outgroup; red: *M. m. musculus*, blue: *M. m. domesticus*; the numbers are bootstrap support based on 1000 replicates.
